# Long Term Virucidal Activity of Rosin Soap on Surfaces

**DOI:** 10.1101/2022.04.29.490117

**Authors:** Stephen H Bell, Derek J Fairley, Hannele Kettunen, Juhani Vuorenmaa, Juha Orte, Connor G G Bamford, John W McGrath

## Abstract

Microbiocidal products for decontaminating both animate and inanimate surfaces that may act as vectors for disease transmission are a well-established method for inactivating viruses of clinical significance. There are a wide variety of such microbiocidal products that can be purchased commercially, many with different active ingredients. We have recently shown that Rosin soap (derived from Tall Oil that has been produced by trees) is a highly effective virucidal product against enveloped viruses like influenza A virus and SARS-CoV-2 when tested in solution. Here we further demonstrate that Rosin soap retains its virucidal activity against influenza A virus and SARS-CoV-2 when applied to surfaces, such as plastic, glass and steel, either before or after virus inoculation. The virucidal activity extended out to seven days post administration. Together, our results show the potential for Rosin soap-based disinfectants to be used as a surface coating to protect against clinically relevant enveloped viruses, such as influenza viruses and coronaviruses.

## Introduction

There exists a wealth of viruses that can spread between humans, causing disease, and resulting in significant global morbidity and mortality. One prescient example is SARS-CoV-2 which has led to >450 million cases and contributed to >6 million deaths as of March 2022 (1). Respiratory viruses like SARS-CoV-2 (alongside other coronaviruses), Influenza and Respiratory Syncytial Virus can infect either via aerosol exposure, or through direct contact with contaminated droplets on surfaces (2): such viruses are known to persist and remain viable on surfaces for up to 3 days (2, 3). Recent data has also shown that the Omicron SARS-CoV-2 variant is more stable on surfaces such as glass, stainless steel, and polypropylene than earlier variants (4). As such surface contamination in, for example, homes, hospitals, communal areas, airports, and public transport, can act as both a source and a sink for viral transmission (2, 3, 5, 6).

The current COVID-19 pandemic has accelerated the demand for both hand and surface sanitising products, with surface disinfectants providing an effective strategy for mitigating infections (7, 8). Whilst many commercial viral sanitising products inactivate viruses upon contact, few retain their virucidal activity when dried onto surfaces. As such frequent re-application is required. Development of a surface sanitising product which retains its efficacy when dried onto a surface would thus have commercial application both in terms of enhanced public health protection and cost saving (by negating the need for continual application) (9, 10).

We have recently shown that rosin (Tall Oil) can act as a potent viricide when in suspension (11). Rosin is a natural substance secreted by coniferous trees as a defense mechanism against wounds in tree bark. It is a complex mixture of sodium salts and acids including, abietic acid, dehydroabietic acid, pimaric acid and palustric acid which may be a contributing factor to its antibacterial and antiviral properties (11, 12, 13). In our previous study incubation of either Influenza A Virus (IAV), RSV or SARS-CoV-2 with a 2.5% w/v solution of Rosin soap resulted in a rapid and potent (1,000,000-fold reduction) in virus titre within 5 minutes (11). Inactivating viruses when in suspension has however been shown to be less difficult than when they dried onto a surface (14). We now demonstrate that Rosin soap also reduces the infectivity of IAV on a diverse range of surfaces, with the treated surface retaining virucidal activity for up to 7 days.

## Materials and Methods

### Cell culture

In order to grow and assess the infectivity of viruses in this study, mammalian cell lines (MDCK; Madin-Darby Canine Kidney) were cultured in DMEM (high glucose) supplemented with foetal bovine serum (v/v 5%). Cell cultures were maintained in flasks (T175cm^2^) and passaged routinely as described in Bell et al., (11).

### Rosin soap

Rosin soap used in our assays was produced from crude Tall Oil by Forchem Ltd (Rauma, Finland). The Rosin soap is a water-based solution obtained from dried Rosin salt, which consists of less than 10% sodium salts of Tall Oil fatty acids and over 90% sodium salts of resin acids (11). Resin acids (abietic acid, dehydroabietic acid, pimaric acid and palustric acid) and fatty acids in the product originate from the coniferous trees *Pinus sylvestris* L. and *Picea abies* L.

### Viruses

Stocks of influenza A virus (WSN) were prepared using standard virology techniques on MDCK. All viral preparations were carried out in accordance with Bell et al., (1). All virus work was carried out in the Biological Safety Level (BSL) 2 facilities at Queen’s University Belfast, Belfast, UK.

### Surfaces

Three surfaces were used to assess Rosin soap efficacy: plastic, glass, and steel. A plastic petri dish (Sartedt), glass petri dish (Greiner) and stainless-steel plate (EsportsMJJ; 1mm x 100mm x 100mm) were used in the assays. Stainless steel is designated in the British Standard, BS EN 16777:2018 as a non-porous test surface for disinfectants.

### Dried treatment assay of virus infectivity

150μl (2.5% w/v) of treatment (2.5% w/v Rosin soap powder) was dried onto the glass, plastic or steel surfaces. 50μl of virus (concentration, 1× 10^8^ Tcid50/mL) was applied onto the dried treatment (10mL of virus was concentrated using centrifugation through Amicon® Ultra-15 Centrifugal Filter Units (Merck). After 5 minutes at room temperature 150μl of DMEM (high glucose) media was added and pipette mixed ten times. 10-fold dilutions were carried out in a 96 well plate to which 100μl of DMEM (high glucose) supplemented with foetal bovine serum (v/v 5%) was added to each well. Following dilution of the virus, permissive cells were added (100μl) and incubated for between 2-3 days at 37°C.

### Dried virus assay of virus infectivity

50μl of virus (concentration, 1× 10^8^ Tcid50/mL) was dried onto different surfaces (glass, plastic and steel). 10mL of virus was concentrated using centrifugation through Amicon® Ultra-15 Centrifugal Filter Units (Merck). Efficacy against steel was assessed at room temperature and 4°C. 150μl treatment (2.5% w/v Rosin soap powder) was applied to the dried virus. After 5 minutes at room temperature 50μl of DMEM (high glucose) media was added. 10-fold dilutions were carried out on a 96 well plate to which 100μl of DMEM supplemented with foetal bovine serum (v/v 5%) was added to each well. Following dilution of the virus, permissive cells were added (100μl) and incubated for between 2-3 days at 37°C.

## Results

### Efficacy of Rosin soap on plastic surfaces

We have shown previously that Rosin soap (2.5% w/v) has been shown to reduce infectivity of the both enveloped virus IAV (WSN strain) and SARS-CoV-2 after 5 mins incubation (11). As these assays were carried out in solution, we now have further assessed the efficacy of Rosin soap on surfaces, where either the inoculum or the product is dried on to commonly occurring surfaces.

In the first instance, IAV was dried onto a plastic surface substrate to simulate viral contamination of a surface. To this end, IAV stock was added to the plastic surface and allowed to airdry (full dryness was observed after 1 hour) after which Rosin soap (2.5% w/v) was applied. After 5 minutes incubation, residual infectivity was assessed by harvesting virus from the surface through application of cell culture media and assessing infectivity. The addition of liquid Rosin soap (2.5% w/v) to dried IAV reduced the amount of virus to undetectable levels when at room temperature for 5 minutes (Figure 1A). The addition of controls such as, virus in suspension (DMEM; high glucose) and dried virus without the exposure to Rosin soap (2.5% w/v) allowed for viral loss during the drying phase to be accounted for. Drying IAV did not cause it to be inactivated, it was only after the addition of Rosin soap (2.5% w/v) that it was inactivated, dropping 10,000-fold (Figure 1A).

**Figure 1.**
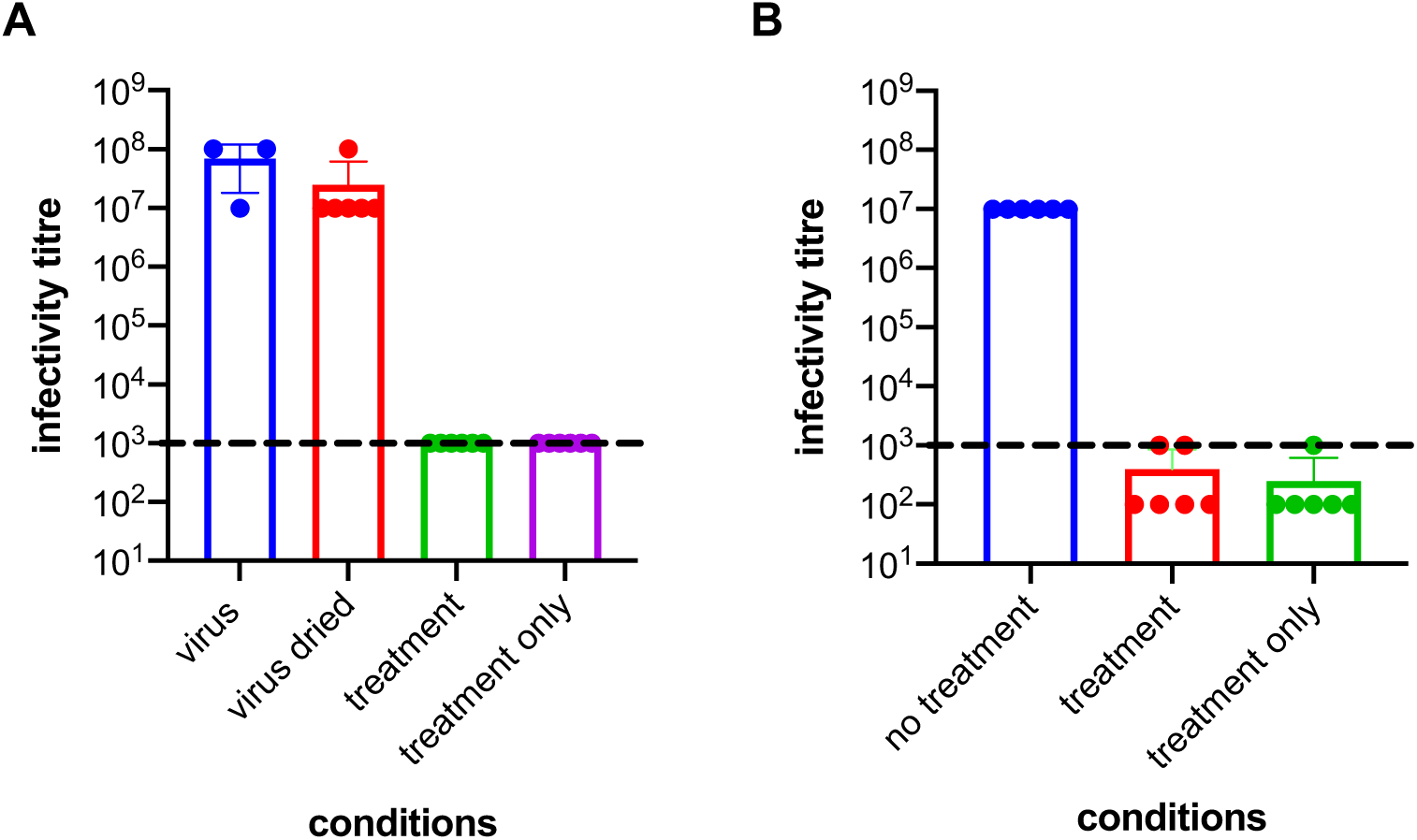
Effect of dried Rosin soap on IAV and Rosin soap in solution on IAV surface infectivity. For Figure 1A) virus (virus levels in stock solution), virus dried (virus levels when dried on plastic), treatment (virus exposed to treatment) and treatment only (treatment in solution, not exposed to virus). For Figure 1B) no treatment (virus in solution, not exposed to treatment), treatment (virus in solution, exposed to dried treatment) and treatment only (dried treatment, not exposed to virus). 1A) Dried IAV was exposed to Rosin soap (2.5% w/v) for 5 minutes. Controls for this assay included dried virus and virus in solution (DMEM (high glucose) supplemented with foetal bovine serum (v/v 5%)) to account for losses that may occur during the drying process. A treatment only (Rosin soap (2.5% w/v)) control was also added. 1B) Dried Rosin soap (2.5% w/v) was exposed to IAV for 5 minutes. A no treatment (Rosin soap 2.5% w/v) control was added to determine the reduction in virus levels post exposure to dried Rosin soap (2.5% w/v). Dashed lines denote the dilution at which Rosin soap is toxic to MDCK cells.

To investigate if Rosin when applied onto a surface retained its viricidal activity we dried Rosin soap (2.5% w/v) to the plastic substrate and left the rosin solution to air dry for 1 hour (until dry). IAV was subsequently added to the Rosin soap coated surface and incubated for 5 minutes at room temperature. At the end of this incubation period, the viral inoculum was harvested, and residual infectivity assessed. Rosin soap (2.5% w/v) when dried onto a plastic surface was effective against IAV (WSN) at room temperature for 5 minutes, reducing virus levels by 10,000-fold (Figure 1B). Similar results were obtained for SARS-CoV-2 D614G, with virus levels reduced by 100,000-fold (results not shown). The presence of a no treatment control (IAV virus that has not been exposed to dried Rosin soap (2.5% w/v)), allows for the efficacy of Rosin soap to be determined when compared to the virus levels post treatment exposure.

### Efficacy of Rosin soap on non-plastic surfaces

Inactivation of IAV was also tested on glass and steel surfaces respectively: stainless steel is designated in the British Standard, BS EN 16777:2018 as a non-porous test surface for disinfectants. Steel was tested at both room temperature and at 4°C [viruses have been shown to be more stable at 4°C and have longer survival on smooth surfaces such as steel, with cold steel in particular known to enhance virus stability (4)]. Rosin soap (2.5% w/v) when dried onto both glass and steel surfaces significantly reduced virus levels by (1,000-fold and 100,000-fold respectively) at room temperature for 5 minutes (Figure 2, C to F). At 4°C on steel, 2.5% (w/v) Rosin soap reduced virus levels by (1,000-fold). Liquid rosin soap (2.5% w/v) applied to virus dried onto the surfaces also reduced virus levels at room temperature by (100-fold and 1,000-fold). Liquid Rosin soap (2.5% w/v) was also effective against dried virus when applied at 4°C for 5 minutes on steel; virus levels were reduced by 1,000-fold (Figure 2, G and H). These assays highlight that Rosin soap activity when either directly dried onto a surface or applied to a dry virus is independent of surface composition.

**Figure 2.**
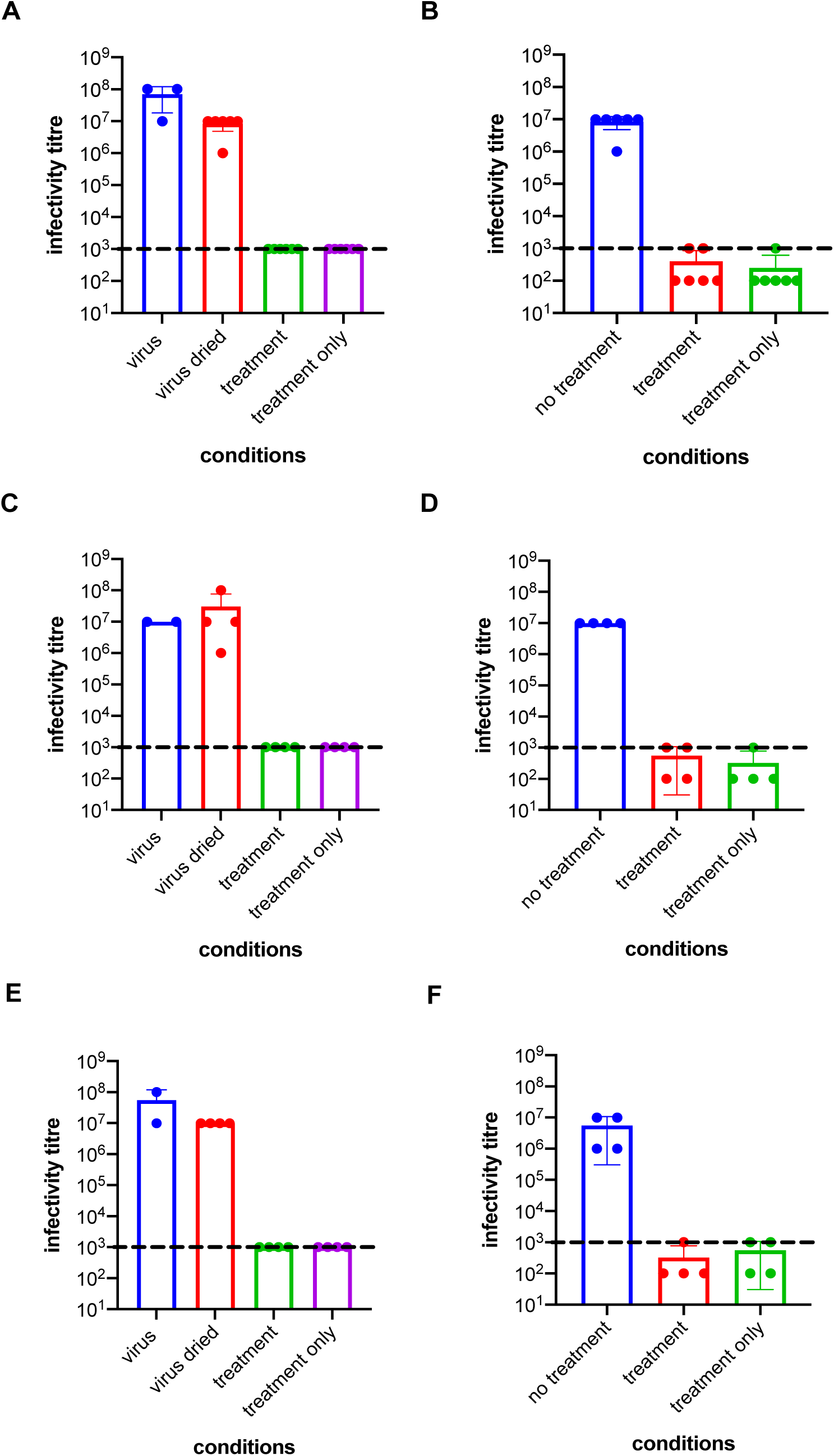
Effect of Rosin soap on IAV infectivity after virus was dried to a glass and steel (room temperature and 4oC) surface. For Figure 2 A, C and E) virus (virus levels in stock solution), virus dried (virus levels when dried on plastic), treatment (virus exposed to treatment) and treatment only (treatment in solution, not exposed to virus). For Figure 2 B, D and F) no treatment (virus in solution, not exposed to treatment), treatment (virus in solution, exposed to dried treatment) and treatment only (dried treatment, not exposed to virus). A) IAV was dried on a glass surface before liquid Rosin soap (2.5% w/v) was applied. B) Rosin soap (2.5%) was dried on a glass surface and suspended IAV (DMEM; high glucose supplemented with foetal bovine serum (v/v 5%)) was applied on top of the dried Rosin. C) IAV was dried on a steel surface (room temperature) before liquid Rosin soap (2.5% w/v) was applied. D) Rosin soap (2.5%) was dried on a steel surface and suspended IAV (DMEM; high glucose supplemented with foetal bovine serum (v/v 5%)) was applied on top of the dried Rosin. E) IAV was dried on a steel surface (4°C) before liquid Rosin soap (2.5% w/v) was applied. F) Rosin soap (2.5%) was dried on a steel surface (4°C) and suspended IAV (DMEM; high glucose supplemented with foetal bovine serum (v/v 5%)) was applied on top of the dried Rosin. All of the assays were carried out at room temperature with the exception of E and F at 4°C. Assays were carried out in two replicates in three independent experiments. A, C and E include virus only (IAV in suspension (DMEM; high glucose supplemented with foetal bovine serum (v/v 5%)), dried virus and treatment (Rosin soap 2.5% w/v) only controls. These controls allow for any viral loss during the drying process to be accounted for as well as the efficacy of Rosin soap when conpared to intial virus levels. B, D and F include, no treatment controls consisting of suspended IAV (DMEM; high glucose supplemented with foetal bovine serum (v/v 5%)) and a treatment only control (Rosin soap 2.5% w/v). This allows pre and post treatment virus levels to be determined. Dashed lines denote the dilution at which Rosin soap is toxic to MDCK cells.

**Figure 3.**
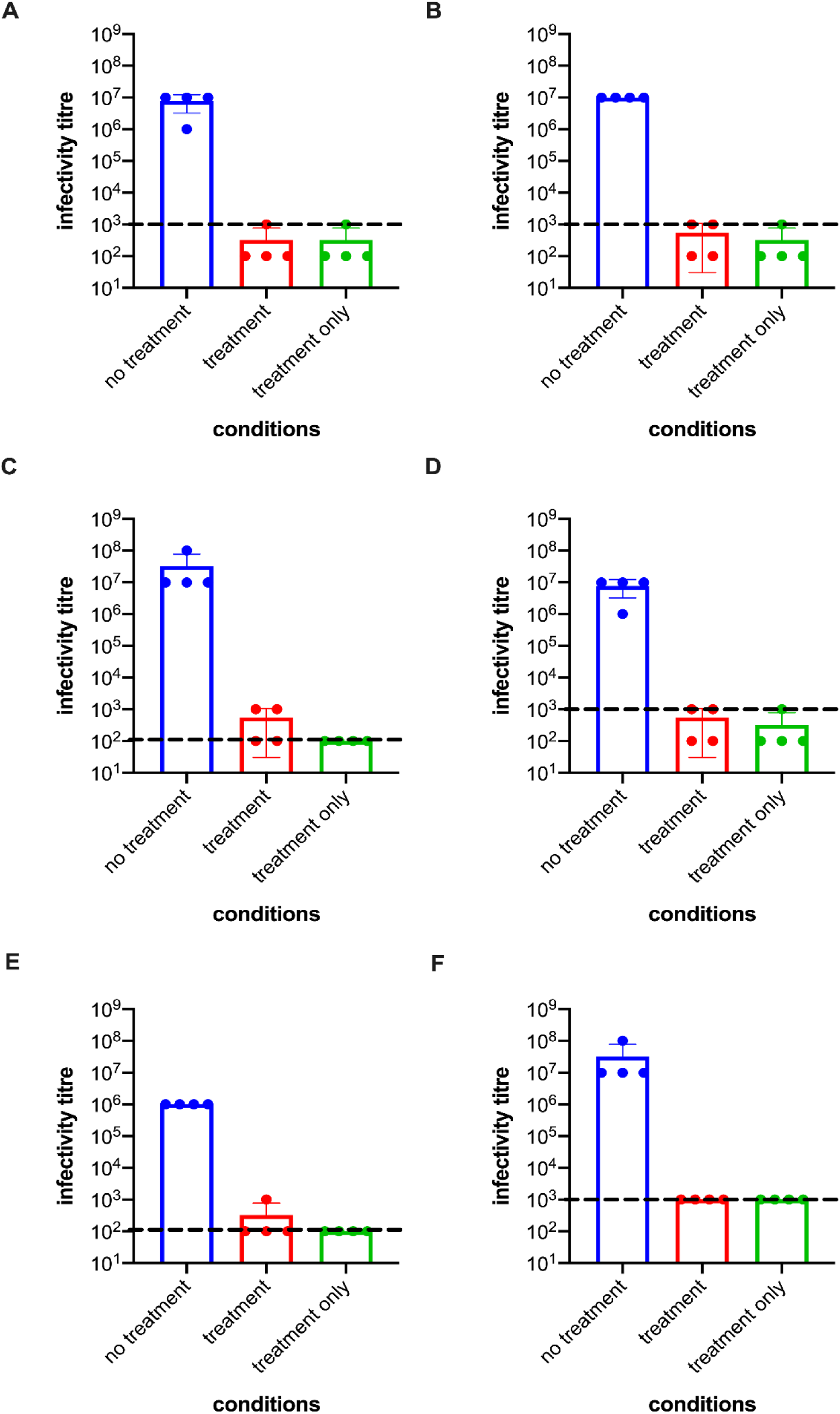
Efficacy of Rosin soap on IAV infectivity after 24 hours and 7 days. For Figure 3)no treatment (virus in solution, not exposed to treatment), treatment (virus in solution, exposed to dried treatment) and treatment only (dried treatment, not exposed to virus). Rosin soap (2.5% w/v) was dried to plastic, glass, and steel at 4°C (A, C and E respectively) for 24 hours and 7 days (B, D and F respectively) before exposure to IAV. All assays include, no treatment controls consisting of suspended IAV (DMEM; high glucose supplemented with foetal bovine serum (v/v 5%)) and a treatment only control (Rosin soap 2.5% w/v). This allows pre and post treatment virus levels to be determined. Dashed lines denote the dilution at which Rosin soap is toxic to MDCK cells.

### Long term efficacy of Rosin soap on surfaces

Protection of surfaces relies on the presence of adequate levels of active product. To assess the longevity of Rosin soap (2.5% w/v) efficacy it was dried on surfaces (plastic, glass, and steel (steel was assessed at room temperature and 4°C) and left for either 24 hours or 7 days before inoculation with IAV virus for 5 minutes at room temperature. Rosin soap (2.5% w/v) retained its efficacy after 24 hours and 7 days respectively. When dried on plastic, glass, and steel (4°C) virus levels after 24 hours were reduced 1,000-fold, 10,000-fold, and 100-fold respectively. After 7 days, efficacy against IAV on each for the three surfaces (plastic, glass, and steel (4°C) gave rise to a 10,000-fold, 100-fold, and 1,000-fold reduction in virus titre respectively.

## Discussion

Pathogenic viruses can be spread via surfaces, such as animate substrates like skin, or inanimate surfaces like plastics with the highest surface levels of virus usually found directly after exposure (3, 15, 16). Surface decontamination is thus a useful public health strategy for preventing the transmission of viruses of clinical significance (2, 7, 16). We have previously demonstrated that Rosin soap has virucidal activity against enveloped viruses like IAV and SARS-CoV when tested in solution (11). We now build on this previous research and show that Rosin soap (2.5% w/v) has long-term virucidal activity against IAV and SARS-CoV when dried onto surfaces.

Rosin soap is naturally occurring product composed of Tall Oil fatty acid sodium salts and resin acid sodium salts. When dried, we hypothesised that water would evaporate and leave a residue composed of those sodium salts and retain antiviral efficacy. It is important to understand if, when natural products such as Rosin soap are dried onto a surface that they retain their antiviral efficacy if a commercial product is to be produced. Many antiviral assays have been carried out using natural products such as caffeine, ascorbic acid, and essential oils (17, 18, 19). However, their activity when dried has not been determined.

The main finding of our work is the long-term efficacy of Rosin soap when dried onto surfaces, with the product significantly reducing (by 100,000-fold) the concentrations of both IAV and SARS-CoV-2 D614G. Furthermore, using IAV as the model test virus, Rosin soap retained its efficacy on all surfaces tested (plastic, glass, and steel) even after 7 days. This virucidal activity against IAV and SARS-CoV-2 D614G is consistent with that seen for Rosin soap in suspension (11). As previously noted by Bell et al., (11) the cytotoxicity of Rosin soap to *in vitro* cell lines is such that we cannot determine if virus has been reduced to undetectable levels, i.e., complete inactivation.

Previous studies have shown the efficacy of natural products against enveloped viruses such as IAV and SARS-CoV-2, with both liquid chalk and dried copper-based compound coatings retaining virucidal activity after drying - the latter being effective for up to 2 hours (20, 21). In contrast, dried Rosin soap remained active throughout the 7 days of this study with its efficacy demonstrated on multiple surfaces that are found frequently within homes and communal areas (plastic, glass, and steel). As surfaces are a well-known source of virus infection, where persistence remains for prolonged periods of time (3, 7, 22), the long-term efficacy of Rosin soap is significant and commercially important, as it negates the need for continual reapplication to those surfaces (5, 10). As viruses can persist and are often transmitted on surfaces commonly found in high touch areas such as plastic, glass and steel (3, 4), protecting such areas with a surface-active coating, that has long term efficacy, such as Rosin soap would have obvious public health advantages.

## Acknowledgements

This work was funded by support from Hankkija Oy and Forchem Oy.

